# Human retina trades single-photon detection for high-fidelity contrast encoding

**DOI:** 10.1101/2022.12.15.520020

**Authors:** Markku Kilpeläinen, Johan Westö, Anton Laihi, Daisuke Takeshita, Fred Rieke, Petri Ala-Laurila

**Affiliations:** Department of Psychology and Logopedics, University of Helsinki, Helsinki, Finland; Molecular and Integrative Biosciences Research Programme, University of Helsinki, Helsinki, Finland; Department of Physiology and Biophysics, University of Washington, Seattle, WA; Neuroscience and Biomedical Engineering., Aalto University, Espoo, Finland

## Abstract

We lack a fundamental understanding of how the spike output of the retina enables human visual perception. Here we show that human vision at its ultimate sensitivity limit depends on the spike output of ON but not OFF parasol (magnocellular) ganglion cells. Surprisingly, nonlinear signal processing in the retinal ON pathway precludes perceptual detection of single photons in darkness, but enables quantal-resolution discrimination of differences in light intensity.

---

Detection of the dimmest lights offers perhaps the most remarkable example in biology of how sensory performance can approach the fundamental limits of physics. Classic psychophysical experiments showed that dark-adapted humans can detect less than a dozen light quanta^1–3^. Despite decades of work, the neural mechanisms that define this remarkable performance remain poorly understood. A key unresolved question is whether there is a neural threshold (nonlinearity) somewhere in the visual pathway requiring a minimum number of absorbed quanta to discern signals from neural noise. Current theory (“classical model”, see Fig. 1a) proposes that signals initiated by each photon absorption propagate linearly or near-linearly through visual circuits in the retina before encountering a perceptual decision criterion or threshold in the brain. In this model, perception, with an appropriate criterion, can access signals produced by each individual photon^4,5^. However, past psychophysics experiments do not unambiguously support this classical model^6–8^.

**Fig. 1.**
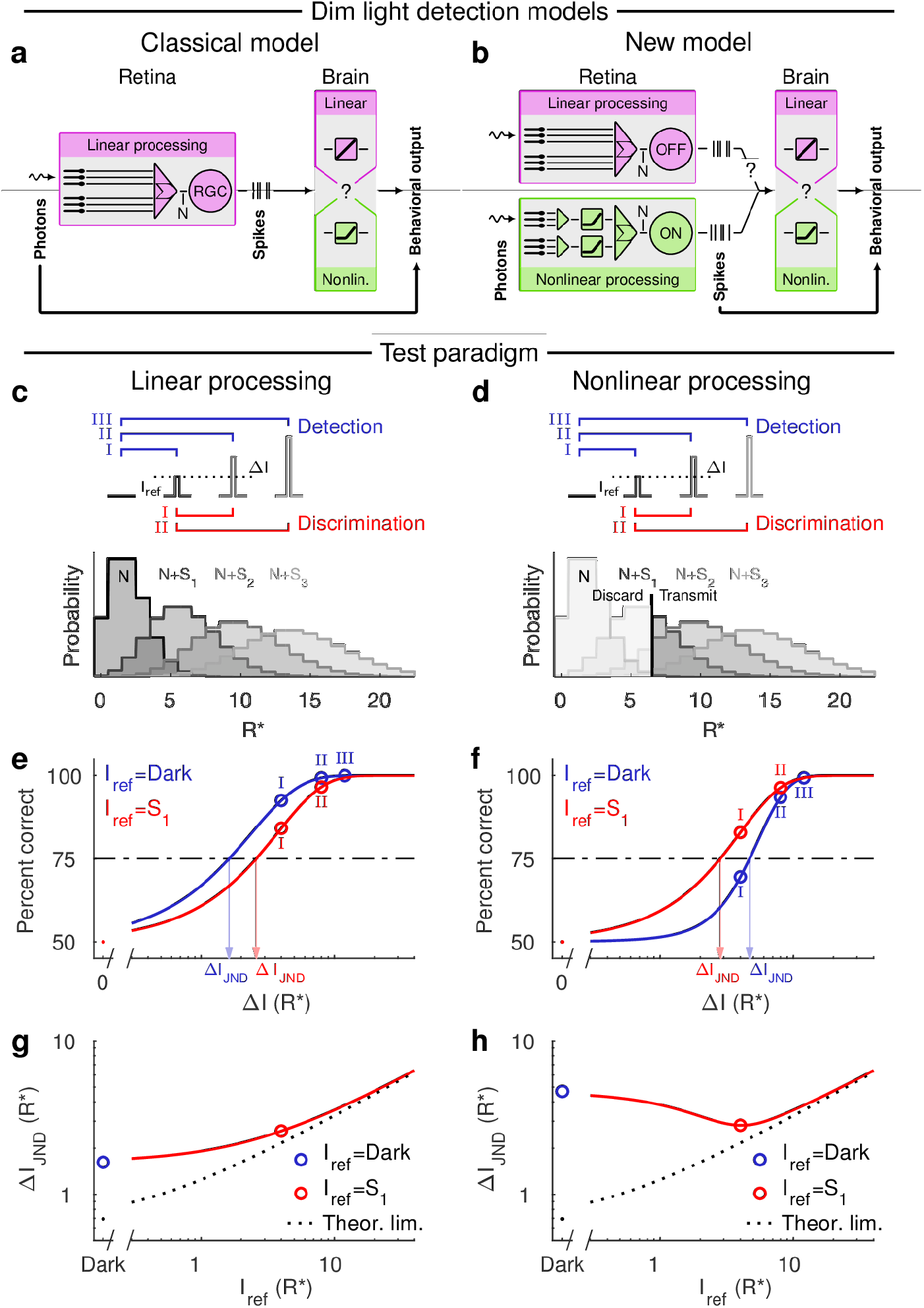
Linear and nonlinear dim light detection models exhibit fundamentally different performance characteristics for detection and discrimination tasks. **a,** The classical dim light detection model assumes linear retinal processing and that behavioral sensitivity is only limited by noise and potentially a downstream thresholding mechanism beyond the retina. **b,** The new model, in turn, postulates that behavioral sensitivity instead is fundamentally limited by a thresholding mechanism along the ON pathway in the retina. **c** and **d**, The presence of thresholding nonlinearities can be observed by evaluating the dimmest light increment needed for detection (blue) and discrimination (red) of dim lights in a two-alternative forced choice (2AFC) setting. Linear processing (**c**) allows an ideal observer to access all isomerization events occurring (R*; Poisson distributed signals S_1_, S_2_, and S_3_, and noise N), whereas a thresholding nonlinearity (**d**) restricts access to only those events that surpasses the threshold. **e** and **f,** Ideal 2AFC performance for detection (blue) of a dim flash against a background noise (N), and for discriminating (red) a probe flash from a refence flash (S_1_). For linear processing (**e**), the detection task is easier than the discrimination task (quantified by the just noticeable difference; ΔI_JND_), whereas the situation is reversed for nonlinear processing (**f**; the blue curve is now on the right side of the red one). **g** and **h,** The JND as a function of the reference flash intensity: the detection task (blue dot) thus corresponds to blank reference flash. The dotted black line shows the theoretical performance limit set by quantum fluctuations in the number of stimulus photons.

Two recent findings challenge the classical model: First, signals traversing the ON (but not OFF) pathway in the primate retina are subject to a thresholding nonlinearity that discards noise and most single-photon responses while transmitting signals originating from coincidence of two or more quanta in a pool of ~1000 rods^9^. Second, recent work in mice shows that behavioral detection of dim flashes in darkness relies on ON RGCs even when the OFF RGCs would allow higher sensitivity^10^. Given the conserved rod bipolar circuit that conveys single-photon signals across mammalian retinae, these two findings suggest a new model in which perception relies on retinal outputs originating in the ON pathway, and those signals are shaped by a thresholding nonlinearity that limits access to individual single-photon responses (see Fig. 1b).

Here, we describe two new results that distinguish between these models. First, we measured the output signals of the primate retina as a proxy for human retinal outputs in stimulus conditions closely matched to human psychophysical experiments. Second, we used a richer perceptual task that separates the models more clearly than a simple detection task alone. Specifically, the two models make different performance predictions for detection and discrimination tasks in a two-alternative forced choice (2AFC) setting. In the detection task, an observer reports which of two stimulus intervals contained a flash. In the discrimination task, the observer reports which of two sequential flashes appears brighter.

Figs. 1c-h show predicted 2AFC task-performance for linear and nonlinear models. Fig. 1c shows the response (signal + noise) distributions for three flash strengths assuming linear processing. Fig. 1d applies a threshold to these distributions. For a linear system, detection of a flash in darkness (blue trace, Fig. 1e) is always easier than discrimination between two flashes with a corresponding difference in flash strength (red trace, Fig. 1e). This is because Poisson distributions arising initially from the photon statistics of light get wider and their overlap increases as the flash strength increases (Fig. 1c). Elimination of small responses by nonlinear processing, however, can lower sensitivity in the detection task without affecting the ability to discriminate between supra-threshold responses. This can permit discrimination of a difference in strength between two dim flashes that is smaller than the detection threshold (Fig. 1f). This phenomenon creates a “dip” when the increment in flash strength (i.e. *I*_probe_ – *I*_ref_) needed to reach the just noticeable difference (JND; 75% correct in our case) is plotted as a function of the intensity of the reference flash (Fig. 1h). Such dips have been reported in contrast discrimination tasks for retinal ganglion cells in guinea pig^11^ and for human psychophysics^12^. This dip does not occur for a linear system with additive or multiplicative noise, where the function instead exhibits a monotonic increase (see Fig. 1g and Supplementary Fig. 3). The dip therefore is an identifying feature of a nonlinear system.

We start by describing the detection and discrimination performance of primate ON and OFF parasol (magnocellular-projecting) RGCs. ON and OFF parasol RGCs are likely of direct relevance for human psychophysics at its sensitivity limit, due to their high contrast sensitivity and abundant rod input (see Ala-Laurila and Rieke^9^). We recorded spiking activity (cell-attached) of dark-adapted ON and OFF parasol RGCs as the retinal rod bipolar pathway outputs (Fig. 2a inset) in response to sequences of dim flashes (Fig. 2a and b). We then used an ideal observer analysis to quantify detection (Fig. 2c, d; blue symbols) and discrimination performance (Fig. 2c, d, red symbols). As in Fig. 1, we defined performance as the light intensity difference (*ΔI*_JND_) needed to distinguish the brighter probe flash from the dimmer reference flash (*I*_ref_) in 75 % of the trials.

**Fig. 2.**
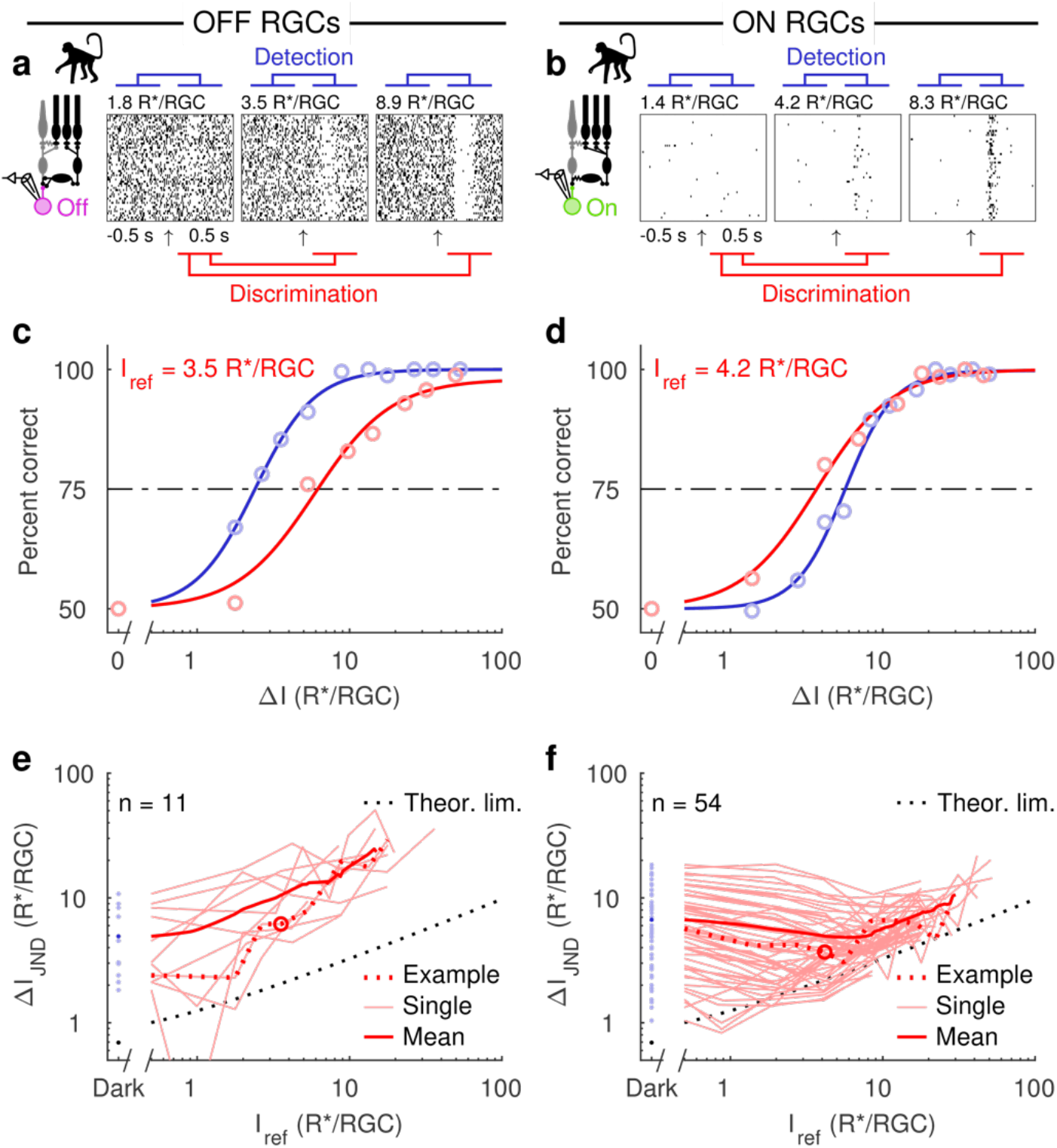
The retinal OFF pathway is linear whereas the ON pathway is nonlinear. **a,b,** Responses of example OFF (**a**, pink) and ON (**b**, green) parasol RGCs as the retinal rod bipolar circuit (black) outputs to a brief dim flash (arrow indicates flash onset). An ideal observer performs the detection or the discrimination tasks by comparing the recorded spike responses. In the detection task, spontaneous activity is compared to flash responses, whereas responses to a probe flash are compared to responses to brighter flashes in the discrimination task. **c,d,** Performance of the OFF (**c**) and ON (**d**) RGCs on the detection task (blue) and on the discrimination task for one probe intensity (red). Markers indicate values computed from measured responses (see Methods), and the continuous lines are fitted Hill functions. The JND is the increase in flash intensity (ΔI) required to reach 75 % correct (dashed line). **e,f,** JNDs as a function of the reference flash intensity (0 for detection) for all OFF (**e**) and ON (**f**) RGCs. The thin lines correspond to individual cells, the dotted lines to the example cells, the thick lines to population averages, and the dotted black line shows the theoretical performance limit set by quantum fluctuations in the number of stimulus photons.

ON and OFF parasol RGC performance differed fundamentally: OFF parasols had better detection performance than discrimination performance, consistent with linear processing (Fig. 2c). ON parasols, in contrast, had better discrimination performance than detection performance (Fig. 2d), consistent with nonlinear processing. Further, the *ΔI*_JND_ as a function of the reference flash intensity rose monotonically for OFF parasol responses and exhibited a clear dip for ON parasol responses (Fig. 2e & f). Strikingly, the ON parasol performance approached the theoretical limit set by Poisson fluctuations of flash-induced visual pigment activations in the discrimination task but not the detection task (dashed black lines in Fig. 2e & f). OFF parasol performance was worse than that of ON parasol cells for discrimination (*P*=0.0007, Welch’s *t*-test, Cohens *d*=2.55) but similar for detection (*P*=0.15, Welch’s *t*-test; Cohen’s *d*=0.41; blue symbols in Fig. 2 e & f, see also Ala-Laurila and Rieke^9^).

How do the differences in detection vs discrimination based on responses of ON and OFF parasol RGCs relate to perception? To answer this question, we evaluated the psychophysical performance of human observers using the same 2AFC tasks. In the detection task, the observer reported whether a dim flash was present in the first or the second of two stimulus intervals (see Detection in Fig. 3a). In the discrimination task, the observer reported which of the two intervals contained a brighter flash (see Discrimination in Fig. 3a). Discrimination performance exceeded detection performance (Fig. 3b, *P*<0.01, Student’s *t*-test, Cohen’s *d*>2, for all observers, see Methods), leading to a clear dip in the *ΔI*_JND_ –function (Fig. 3c). This was the case for all five observers, and the average discrimination performance came strikingly close to the theoretical limit (black dashed line in Fig. 3c), while the detection performance was far from it. Human performance on the discrimination task was thus superior to OFF parasols but slightly inferior to ON parasols (Fig. 3d), supporting the dependence of behavior on responses of ON rather than OFF parasol cells under these conditions and in line with earlier findings in mice^10^.

**Fig. 3.**
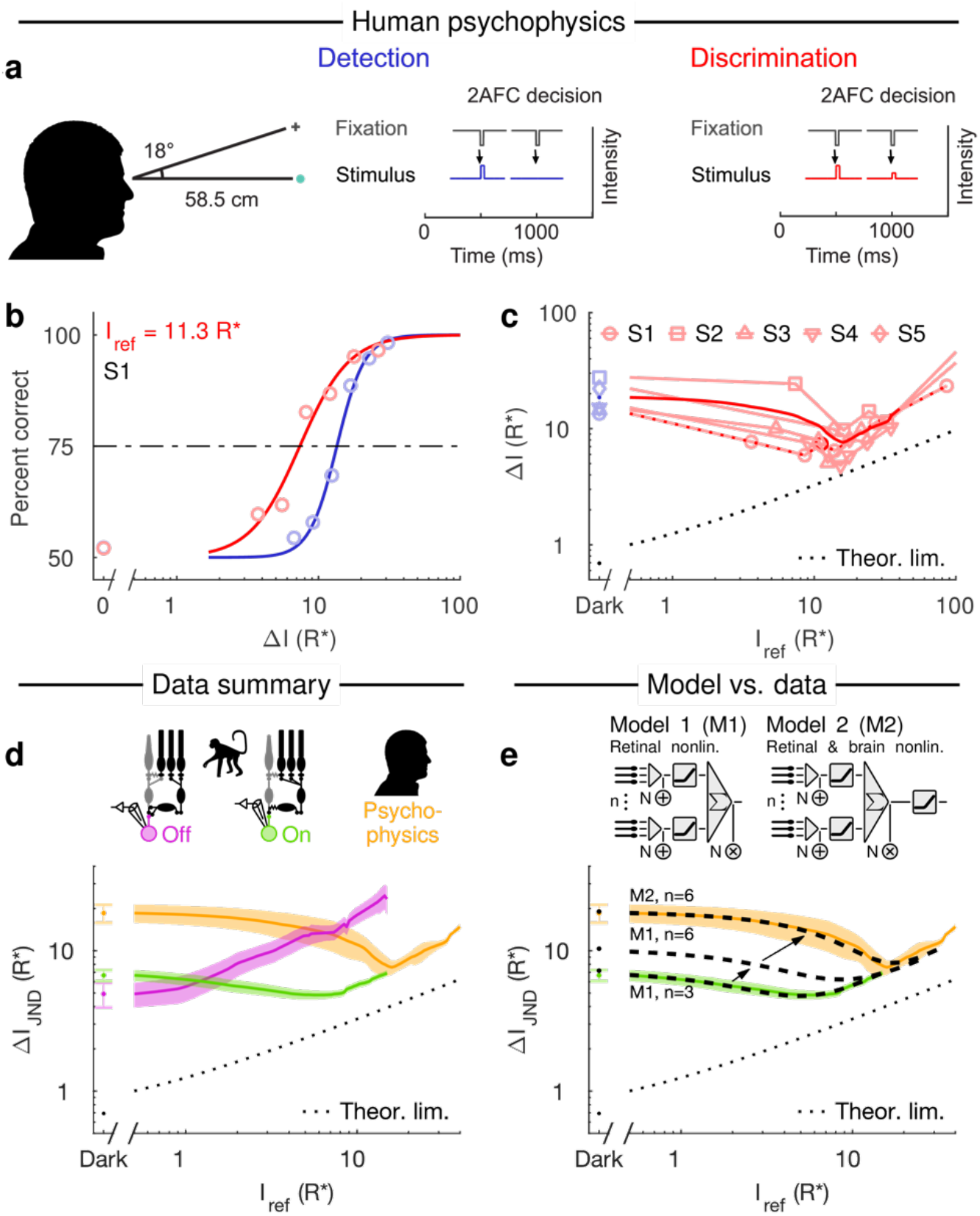
The performance of human observers agrees with the new dim light detection model where performance is fundamentally limited by retinal thresholding nonlinearities along the ON pathway. **a**, The psychophysical experiment. In the detection task, the observer had to report which of two intervals that contained the flash. In the discrimination task, the observer had to report which of two intervals that contained the brighter flash. **b**, Psychometric functions of a representative observer: the detection task (blue) and the discrimination task (red) for one reference flash intensity. **c**, JNDs (75 % correct) as a function of the reference flash intensity (Dark for detection) for all observers. Thin lines correspond to individual observers, the dotted line denotes the example observer in **b**, and the thick line denotes the population average. **d**, Average JNDs as a function of the reference flash intensity for human observer (orange), OFF RGCs (magenta), and ON RGCs (green). **e**, Model predictions for ON RGCs (M1, *n* = 3, *θ* = 2; similar to the parameters used in Ala-Laurila and Rieke^9^), an ON RGC model with twice the number of subunits (M1, *n* = 6, *θ =* 2), and for a model with two thresholding nonlinearities (M2, *n* = 6, *θ =* 2): one in the retina and one downstream of the retina. The dotted black line shows the theoretical performance limit set by quantum fluctuations in the number of stimulus photons.

Human behavior and ON parasol RGCs exhibit nonlinear processing. However, the dip in the *ΔI*_JND_ –function for human behavior is located at higher intensities than that for ON parasol RGCs. This change in the location of the dip suggests additional thresholding beyond the retina. We tested this hypothesis by comparing two distinct models: one with a post-retinal nonlinearity in the brain (Fig 3e, M2) and one without it (M1). These models were based on ON RGCs because their discrimination performance was better than that of OFF RGCs. The aim was to test whether a simple model could link human psychophysics at the sensitivity limit of vision to retinal RGC outputs. The retinal part of both models is identical, with single-photon signals and noise (additive and multiplicative) being summed and thresholded by retinal subunits corresponding to ON-cone bipolar cells, as in Ala-Laurila and Rieke^9^. The retinal nonlinearity is followed by a second nonlinearity in M2, whereas M1 has a linear readout of the retina. For both models, we first fitted (least squares) the number of subunits (*n*), the additive and multiplicative noise, and the threshold (*θ_1_*) to describe the average performance of the ON parasols (Fig. 3e; M1), leading to *n* = 3 and *θ_1_= 2*, close to earlier estimates^9^. This shows that coincidence of a minimum of two single-photon responses in a subunit of ~2000 rods is required to create the response properties of ON parasol ganglion cells (see Methods). Next, we tested if access to more than a single RGC could explain the difference between behavior and ON parasol responses, since the retinal stimulus size in psychophysics corresponds to ~1.5 times the RGC receptive field. Increasing the number of subunits, as demonstrated by doubling their number (M1, *n* = 6, see Fig. 3e) only shifted the curve upward in M1, but did not increase the dip. However, adding the second nonlinearity as in M2, resulted in an excellent fit to the human behavioral data (Fig. 3e; M1, *n* = 6, *θ_1_*=2, *θ_2_*=2). This result is consistent with additional downstream thresholding of ON parasol signals in the brain at the sensitivity limit of vision.

Our results provide three key insights related to the mechanistic and functional underpinnings of human vision at the lowest light levels. First, similar to previous results in mice^10^, human behavior at absolute threshold relies on retinal output signals provided by ON RGCs. This result shows that retinal ON and OFF pathways carry out distinct functional roles for visually-guided behavior at visual threshold, such that behavior relies on the information encoded by increased firing rates of the ON cells for detection of light increments. It remains to be seen if behavioral detection of the weakest light decrements relies on the retinal OFF pathway, as our recent results on mice suggest^13^. Earlier results on primates at photopic light levels indeed suggest that light increments are encoded by ON RGCs and light decrements by OFF RGCs^14^.

Second, our results answer the decades-old question about whether humans can see a single photon; our results indicate that responses to individual photons interact, providing evidence for the perceptual relevance of detection of coincident photons^3^. Thus, the biological circuit design has been optimized for extraordinary discrimination of small light intensity differences as evidenced by the nonlinear ON pathway as compared to the linear OFF pathway. Even if this nonlinear processing strategy leads to a loss of single-photon responses, this loss is almost fully compensated by the elimination of neural noise. This is evident from the observation that the nonlinear ON pathway and the linear OFF pathway perform almost equally well on a light detection task, despite extra single-photon losses in the ON pathway (see also Ala-Laurila and Rieke^9^). Thus, seeing single photons has not been the central optimization goal during evolution. Instead, human vision approaches the theoretical limit of physics in discriminating dim flash intensities by utilizing a strategy where neural noise is eliminated at the expense of single photons to optimize contrast coding in the retinal ON pathway.

Third, single photon responses are separated from noise by thresholding nonlinearities in retinal circuits. The first synapse of the rod bipolar pathway thresholds rod signals to eliminate neural noise and many single-photon responses^15^. Since individual rods very rarely absorb overlapping photons for just-detectable inputs, this first nonlinearity functions as a linear loss mechanism for single-photon responses that is shared for ON and OFF pathways. The last synapse along the ON pathway, on the other hand, requires a coincidence of two or more single-photon responses within a collection of several thousand rods^9^, and this nonlinearity shapes retinal output signals. We add to this picture here by identifying an additional nonlinearity operating in a similar manner downstream of the retina. Such a post-retinal thresholding mechanism in the brain for single-photon detection is similar to previous findings in relay neurons in the thalamus^16^. Together, these results show that a key neural strategy of high-fidelity coding of sparse signals is implemented by a combination of thresholding mechanisms operating at different levels of convergence of the neural circuits together with a large amount of pooling. It is likely that this general strategy and architecture is in use across other circuits and sensory modalities requiring sensitivity that approaches the limits of physics.

## Supplement

Supplementary figures: S1–S3.

## Acknowledgments

We thank Drs. Kristian Donner, Greg Field, Gabriel Peinado, Greg Schwartz, and Mark Georgeson for excellent comments on the manuscript; Matthew Dunkerley, Sathish Narayanan, and Mark Cafaro for the design of the data acquisition software. Support was provided by the Academy of Finland (296269 to PA-L); the Aalto Brain Centre (J.W.); Svenska kulturfonden (J.W.); The Finnish Society of Sciences and Letters (J.W.); The Ella and Georg Ehrnrooth Foundation (M.K.); and the NIH (EY 028111 to F.R.). The datasets and code generated during the current study are available from the corresponding author on request.

## Author contributions

F.R., J.W., M.K., P.A.-L. designed experiments, F.R. and P.A.-L collected RGC data, M.K., A.L. collected psychophysics data, F.R., P.A.-L., J.W., M.K., D.T. analyzed data, F.R., J.W., M.K. & P.A-L wrote the paper.

## METHODS

### Data collection: retinal ganglion cells

All recordings were from ON or OFF parasol retinal ganglion cells in the primate (Macaca nemestrina and fascicularis) retina (eccentricity > 30°). The stimuli consisted of 20 ms flashes (5 to 12 different intensities and 20 to 140 repetitions per intensity) of a uniform circular light spot (diameter 560 μm), delivered from blue and/or green light-emitting diodes (LEDs; peak wavelength *λ_p_* at 460 nm and 510 nm). The procedures have previously been reported^9^. All experiments were done in accordance with the guidelines for the care and use of animals at the University of Washington

### Data collection: psychophysics

#### Observers

Five observers (three male, two female, age 19–29 years) participated in the study. Observer S1 was one of the authors, the rest were naïve to the purposes of the study and received a small monetary compensation. All observers had normal uncorrected vision. The study was conducted in accordance with the principles of the Declaration of Helsinki and in accordance with the guidelines of the University of Helsinki ethical review board, who also approved the study. The participants signed a written informed consent.

#### Stimuli

The stimulus was a homogeneous circular light spot (diameter 1.17° ≈ 315 μm on the retina; *λ_p_* 500 nm) presented during 20 ms at 18° eccentricity in the lower visual field (superior retina). The fixation stimulus was a dim red (*λ_p_* = 680 nm) cross-hair with a diameter of 0.59° (157.5 μm) and a bar width of 0.098° (26.2 μm). The viewing distance was 58.5 cm, which is close to the expected dark focus distance for young adults^17^.

#### Apparatus

The stimulus was produced by a combination of an LED (AND520HB, *λ_p_* = 466 nm), an interference filter (Edmund Optics, *λ_p_* = 500 nm, FWHM 10 nm), ND filters, a 0.8 mm aperture (to produce a more point-like source), a diffuser, and a circular aperture. The intensity of the LED was adjusted at the beginning of each experimental session with ND filters to produce a suitable stimulus range. During the experimental sessions, the light intensity and the flash duration was set using a National Instruments USB-6343 (National Instruments, Austin, TX, USA) DAQ and a custom LED controller. Non-linearities in the stimulation system were corrected for by measuring the system’s input-output function with a UDT Instruments S471 optometer and a UDT 268R Photodiode (Gamma Scientific, San Diego, CA, USA), and by applying the inverse function to the intensity range. The fixation stimulus was produced by a combination of an LED (AND180HRP), an interference filter (*λ_p_* = 680 nm, FWHM 10 nm) and a cross-hair shaped aperture.

#### Procedure

The threshold for discriminating between two weak flashes or a flash and darkness (detection) was measured with a 2AFC method of constant stimuli procedure. Each trial proceeded as follows (see Fig. 3a): The fixation stimulus blinked (i.e., turned off for 100 ms) four times, with a 500 ms period. A flash was presented at 18° eccentricity on the third and fourth fixation blinks, and the observer’s task was to indicate during which blink the flash was stronger. For each run of trials, one of the flashes always had the same reference intensity (*I*_ref_), whereas the other flashes had intensities within a predetermined range (7 steps), with the lowest intensity always being identical to the reference intensity. Consequently, when the reference intensity was 0, one of the flashes was always blank. The purpose of the four fixation blinks was to reduce temporal uncertainty.

One measurement session lasted approximately 1.5-2 hours. In the beginning, the observer adapted to total darkness for 30 minutes. Then the observer performed about 500 trials at their own pace. The length of the session and the precise number of trials depended on the observer’s pace and on the level of subjectively perceived fatigue. Each data point in Fig. 3a represents 144-228 trials for *I*_ref_ = 0 and 74-140 trials for higher *I*_ref_ values. The condition *I*_ref_ = 0 was recorded twice for each observer: both at the beginning and at the end of the study, so as to reveal possible learning effects. The largest difference between the thresholds from the two measurements was 0.04 log units, which is negligible in comparison to the effect of reference intensity. The data from the two detection measurements has thus been pooled.

### Light intensity conversions

In order to compare stimulus intensities between RGC measurements and psychophysics, stimulus intensities were first converted into photoisomerizations per rod per second (R*/rod/s) and later to photoisomerizations per RGC (R*/RGC) or per retina (R*). *Psychophysics*. For psychophysics data, the initial conversion to R*/rod/s was done as follows. Firstly, the stimulus power (*P*_stim_), measured with a photodiode, was converted to a corneal photon flux density (*F*_cornea_) as:

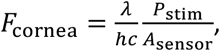

where *λ* is the wavelength of the stimulus light (500 nm), *h* is Planck’s constant, *c* is the speed of light, and *A*_sensor_ is the area of the photodiode (1 cm^2^). Secondly, the corneal photon flux density was converted to a retinal photon flux density (*F*_cornea_) by:

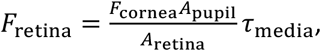

where *A*_pupil_ is the area of the pupil, *A*_retina_ the area of the projected stimulus on the retina (0.077 mm^2^), and *τ*_media_ is the ocular media factor about 45 %^8^. *A*_pupil_ was measured from video recordings of the dark-adapted observers (44.6, 56.4, 32.5, 53.2, and 49.5 mm^2^ for S1, S2, S3, S4, and S5, respectively) while they fixated on the fixation stimulus in darkness (infrared illumination), whereas *A*_retina_ was computed from the stimulus size in visual angles (diameter 1.17°) using the conversion factor from visual angle to retinal subtense at 18° eccentricity (268 μm per °)^18^. Lastly, the photoisomerization rate per rod (*R*_human_) for human observers was obtained as:

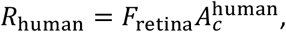

where *A_c_* is the collecting area of rods, obtained from:

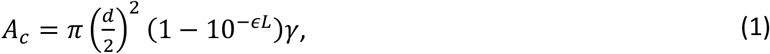

where *d* is the diameter of the rod outer segments (2.27 μm at 18°)^19^, *L* is the length of the rod outer segment (42 μm)^20^, *ϵ* is the specific absorbance (0.019 μm^-1^)^21^, and *γ* is the quantum efficiency of rhodopsin (0.67)^22^. The values above resulted in a collecting area for human rods of 2.28 μm^2^.

#### Stimulus intensity in R*

Flash intensities (*I*) for the psychophysics data are given in R* (per retina). These values were obtained by multiplying the photoisomerization rate per rod (*R*_human_) by the stimulus duration (20 ms) and the total number of rods (10 325) beneath the area covered by the stimulus (rod density of 134 000 rods/mm^2^ at 18°)^19^.

#### Ganglion cell recordings

The intensities used in ganglion cell recordings were converted to photoisomerization rates per rod based on the spectral output of the LEDs, the spectral sensitivity of rods, and a collecting area of 1.40 μm^2^ (eq. 1; *d* = 2 μm and *L* = 25 μm)^22^.

#### Stimulus intensity in R* per RGC

Flash intensities (*I*) for RGCs are given in photoisomerizations per ganglion cell (R*/RGC). These values were determined from the photoisomerization rates (*R*_RGC_) as:

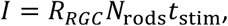

where *N*_rods_ is the number of rods converging on the ganglion cell and *t*_stim_ is the stimulus duration (20 ms). *N*_rods_ was estimated from the size of the spatial RF by multiplying the effective area (integral over the areas of Gaussian weighted annuli up to the size of the stimulus) with the rod density. The size of the spatial RF, in turn, was quantified by presenting dim spots of various sizes for 250 ms and by fitting (least squares) a gaussian shaped RF to map the spot size to observed responses (i.e., assuming a Gaussian weighted linear spatial summation). The size of the spatial RF (2*σ*) was always found to extend beyond the imaged dendritic tree (see Supplementary Fig. 1). We quantified the scaling factor between the size of the spatial RF and the dendritic tree by fitting an ellipse to the edge of the dendritic tree and by comparing the mean axis length of the ellipse to the size of the Gaussian shaped RF. The scaling factor was not significantly different between ON and OFF parasols (*P*=0.36, Welch’s *t*-test; Cohen’s *d*=0.61), and we pooled all data to get a joint mean scaling factor of 2.1. The final estimate of *N*_rods_ was thus obtained by combining the scaling factor, the size of the dendritic tree (diameter: 190 μm at 30°)^23^, the rod density (110 000 rods/mm^2^ at 30°)^24^ and the stimuli size (560 μm). This resulted in a final estimate of 6880 rods per RGC. The given values at 30° are not representative for all recorded RGCs, but the estimate of *N*_rods_ is: the convergence remains fixed as the rod density and dendritic tree size change with increasing eccentricity^25^.

### Data analysis

We quantified discrimination performance in the retina using an ideal observer model that considered both the light response and the tonic firing rate when performing a 2AFC task^9,26^. The recorded spike response was condensed into a single number (*r*) for each epoch by computing the inner product of the binned (10 ms) spike counts with a discriminator over a 500 ms long interval. The discriminator was defined as the mean response difference to the reference intensity (*I*_ref_) and all higher intensities. The ideal observer evaluated the discriminability as a function of flash intensity by comparing the response distribution (the r values from all epochs) to a reference intensity (*I*_ref_) with those obtained at higher intensities (*I*_ref_ + Δ*I*). Performance on the detection task thus corresponded to the special case where *I*_ref_ equaled zero, and for this special case, the responses were calculated from spontaneous activity. The fraction of correct choices by the ideal observer was evaluated as:

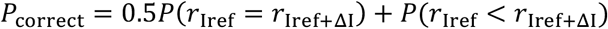

where *r* denotes the response. The just noticeable difference was taken as the increase in flash intensity (Δ*I*) that corresponded to *P*_correct_ = 75 %. This value was obtained by fitting (least squares) a modified sigmoidal Hill function to the data and by determining the intensity where the fit reached 75 % (see Fig. 2c and d). The Hill function was defined as:

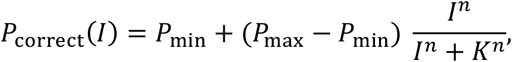

where *P*_min_ is the chance level (= ½), *P*_max_ is the best possible performance ( ≤ 1.0), *I* is the stimulus intensity, *K* is the intensity at 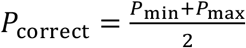, and *n* is the slope. For the ganglion cell data, the number of available data points decreased for each consecutively higher reference intensity, as responses were only available from a fixed number of flash intensities. We solved this by modelling additional response distributions for intermediate flash intensities using normal distributions with the mean and variance fixed to interpolated mean and variance values from the real data (see Supplementary Fig. 2). The modelled response distributions were always interpolated between the reference intensity and the subsequent intensity, and the number of modelled distributions was determined so that the Hill functions could be fit to at least five data points. The psychophysics data was analyzed in an equivalent manner, with the hill functions fit to the response probabilities directly (see Fig. 3b). Ganglion cells were included in the main analyses if they fulfilled a similar sensitivity criteria as established earlier^9^: RGCs had to generate an average spike difference of four to five spikes in response to a brief flash producing 0.001–0.002 R*/rod as compared to the baseline firing rate.

### Modeling

RGC and psychophysics dippers (Fig. 2f and Fig. 3c) were modeled (Fig. 3e) by mapping discrete input events (isomerizations) into discrete outputs (responses) using a subunit model. The stimulus (signal) was distributed uniformly over all subunits and the responses were analyzed in the same way as the recorded data, the only difference being that *P*_correct_ values could be calculated from the modeled output distributions directly. These output distributions were obtained by: 1) pooling (adding) Poisson distributed signals and noise within each subunit, 2) passing the result through a rectifying nonlinearity, 3) summing up the contribution from each subunit, 4) including multiplicative noise, and 5) by thresholding the sum. Summation of the outputs from each subunit was implemented via convolutions, and the multiplicative noise was included by letting the summed subunit output set the mean response for a sequential Poisson process via a gain term (see Field et al.^6^). In total, the model thus had five parameters: the magnitude of the additive noise, the subunit nonlinearity’s threshold, the number of subunits, the gain parameter for the multiplicative noise, and finally the threshold for the final nonlinearity. Optimal values for “free” parameters were in all cases found by minimizing the mean squared error between the model’s dipper function and the target dipper function (from RGC or psychophysics data). The effects of varying each parameter separately is shown in Fig. 3e (number of subunits) and in Supplementary Fig. 3 (remaining parameters).

### Theoretical limit

The theoretical limit (dotted line in figures) correspond to the modeled output when no noise or thresholds are included. That is, it corresponds to linear processing in the absence of noise, and thus reflects JND’s for detection and discrimination due to inherent Poisson fluctuations in the stimulus.

### Statistical analysis

Reported values are means ± s.e.m. MATLAB was used for all statistical analyses, and all reported tests were two-tailed. The statistical significance and effect size of the ON-OFF discrimination threshold difference were determined by conducting a Welch’s *t*-test (and Cohen’s *d*) on the cells’ average discrimination thresholds (averaged over all *I_ref_* values). The statistical significance and effect size of the psychophysics dips were determined separately for each observer by conducting a Student’s *t*-test (with Bonferroni correction) and Cohen’s *d* on the K parameters (75 % correct) of the Hill functions fitted to the detection and discrimination data (solid lines in Fig. 3b). For observers S1, S2, S3, S4, and S5, respectively, the most significant *t*-score was 13.72, 17.98, 4.21, 12.12, 11.19; *P*-values <.001, <.001, .007, <.001, <.001; Cohen’s *d*-values 7.33, 9.61, 2.25, 6.48, 5.98. Pooled SD was used in calculating Cohen’s *d*.

---

**Supplementary Fig. 1.**
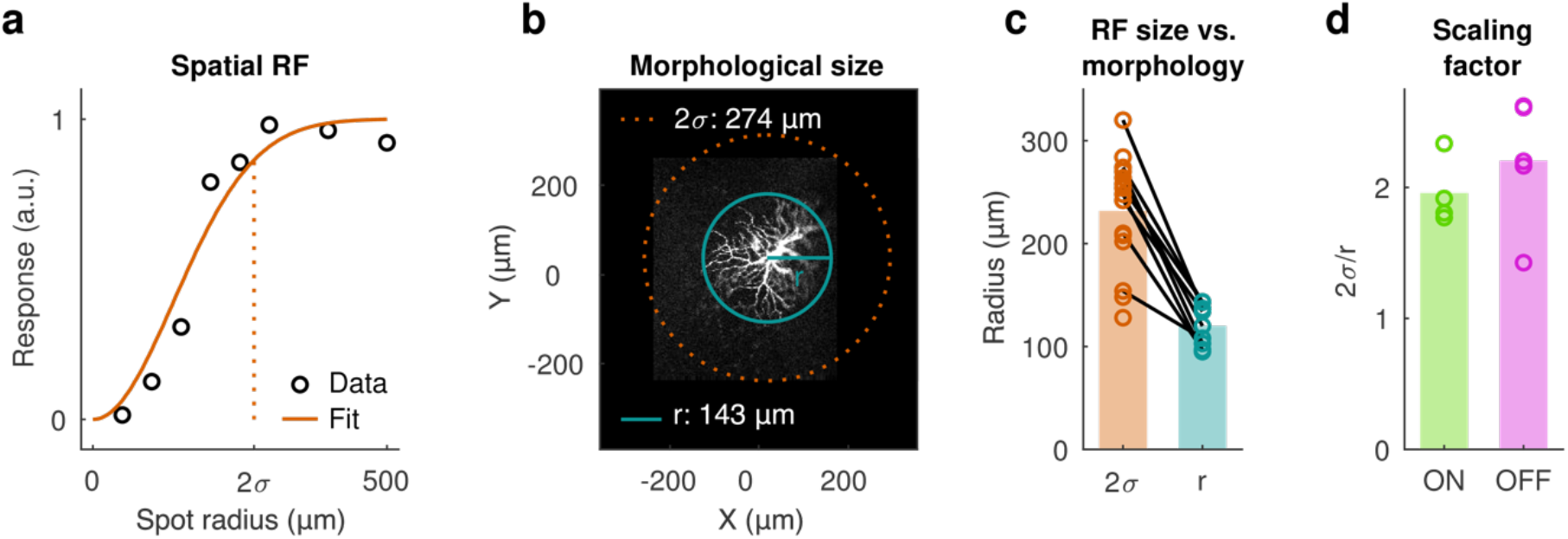
The size of the spatial RF is larger than the width of the dendritic tree in darkness. **a**, The size of the spatial RF was measured by fitting a Gaussian shaped RF to the responses obtained while presenting 250 ms dim spots of various sizes. **b**, The size of the dendritic tree was measured by fitting a circle to the edges of the dendritic tree. **c**, The size of the RF (defined as 2*σ*) was always larger that the dendritic tree (lines denote measurements from the same cell). **d**, The difference could be explained with a scaling factor 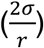 of roughly 2 for both ON and OFF parasols.

**Supplementary Fig. 2.**
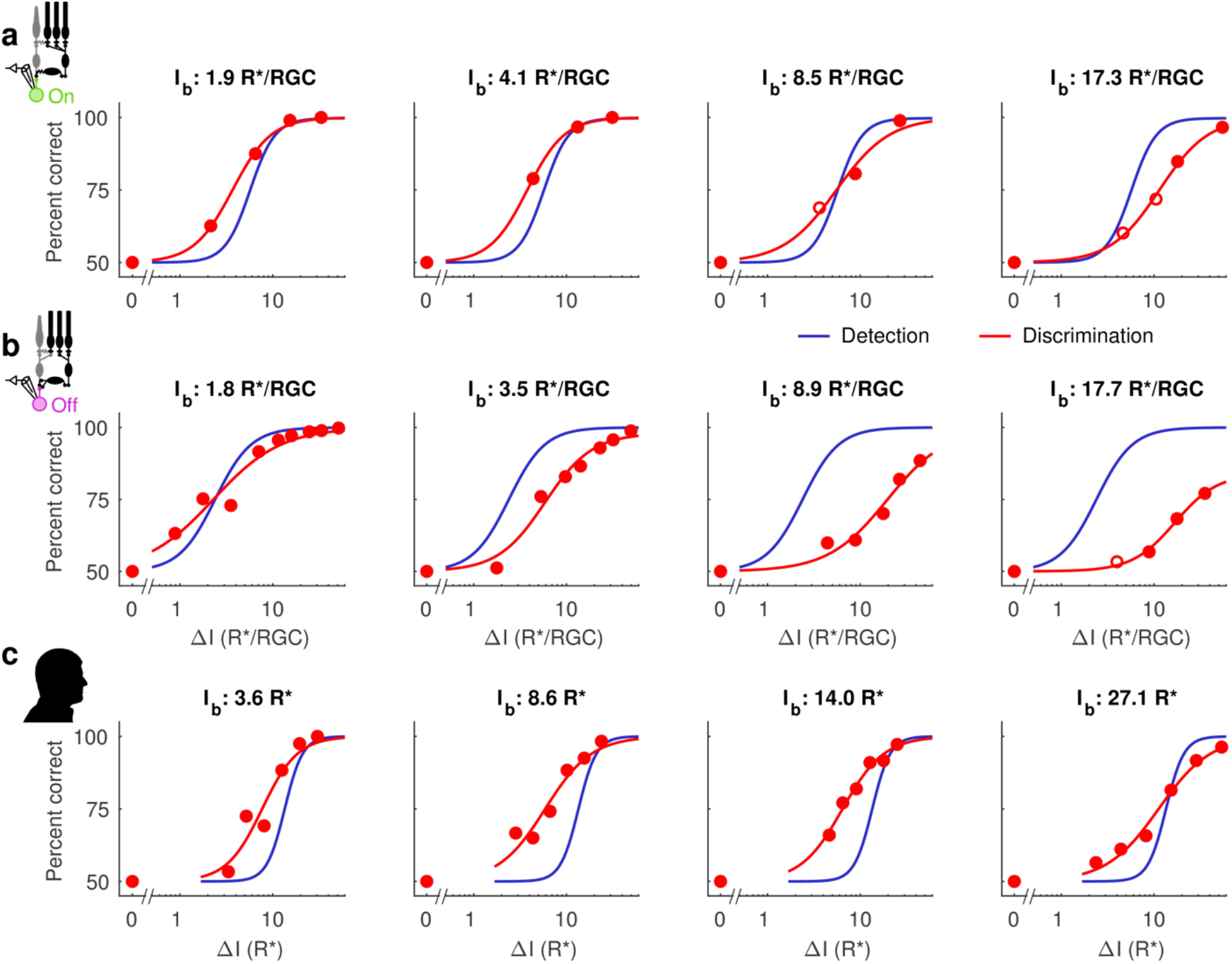
2AFC curves were fitted to all reference intensities and additional response distributions were modelled if needed. **a**, 2AFC curves for four different reference intensities for an example ON RGC. Filled red circles correspond to data points obtained from measured response distributions and open circles correspond to data points obtained from modelled distributions. The red and blue lines denote fitted Hill function for the discrimination and the detection task, respectively. **b**, Same as **a**, but for an example OFF RGC. **c**, Same as **a**, but for one psychophysical observer.

**Supplementary Fig. 3.**
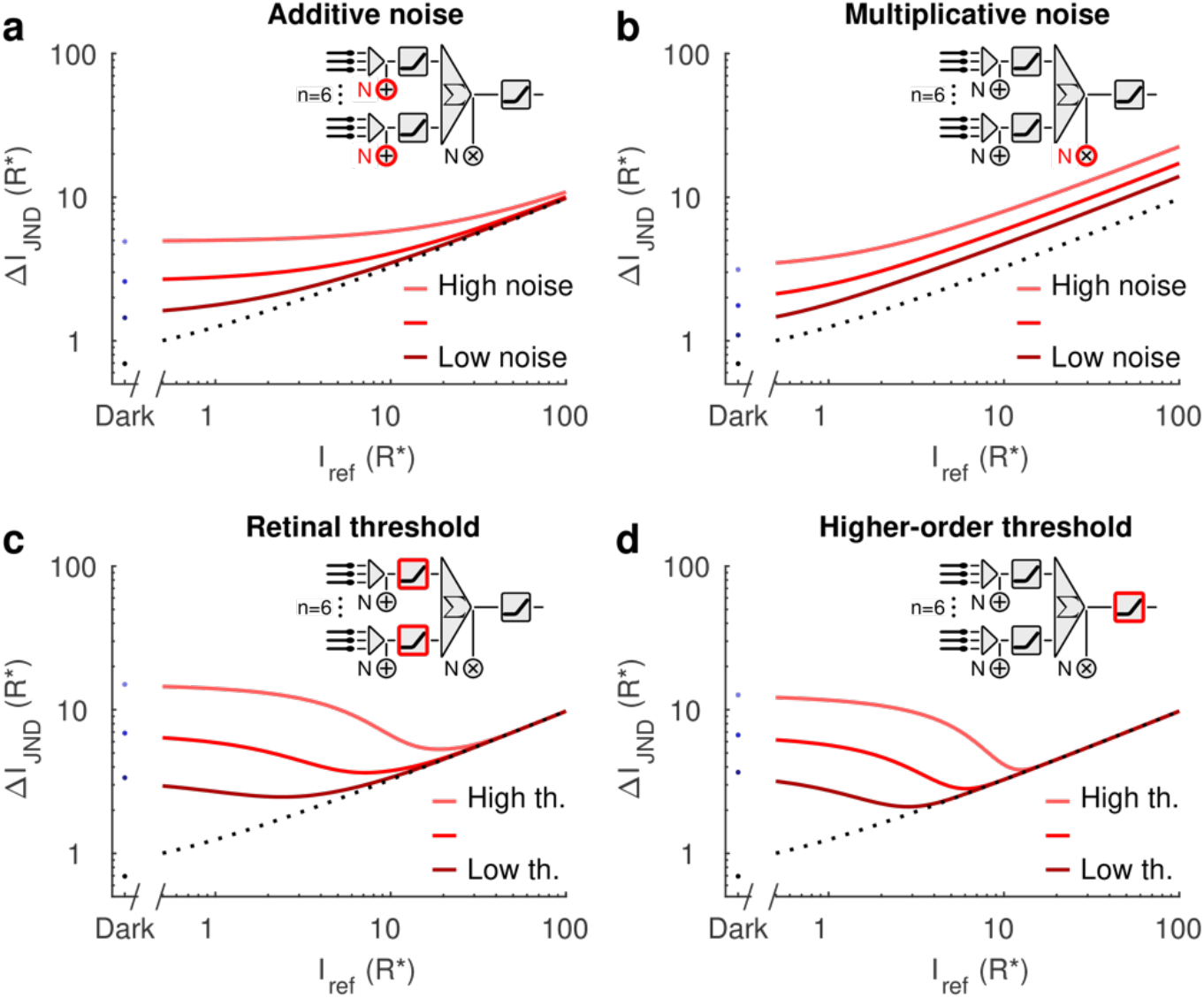
Effects of varying each model parameter separately with respect to the ideal noiseless case without thresholds (dotted black line). **a**, Additive Poisson noise mainly degrades model performance for low reference intensities, and this effect vanishes for larger reference intensities. **b**, Multiplicative noise degrades model performance over all reference intensities. **c** and **d**, Thresholding nonlinearites result in clear “dips” that increase in depth as the threshold increases.

